# A novel interpretable deep transfer learning combining diverse learnable parameters for improved T2D prediction based on single-cell gene regulatory networks

**DOI:** 10.1101/2023.09.07.556481

**Authors:** Sumaya Alghamdi, Turki Turki

## Abstract

Accurate deep learning (DL) models to predict type 2 diabetes (T2D) are concerned not only with targeting the discrimination task but also with learning useful feature representation. However, existing DL tools are far from perfect and do not provide appropriate interpretation as a guideline to explain and promote superior performance in the target task. Therefore, we provide an interpretable approach for our presented deep transfer learning (DTL) models to overcome such drawbacks, working as follows. We utilize several pre-trained models including SEResNet152, and SEResNeXt101. Then, we transfer knowledge from pre-trained models via keeping the weights in the convolutional base (i.e., feature extraction part) while modifying the classification part with the use of Adam optimizer to deal with classifying healthy controls and T2D based on single-cell gene regulatory network (SCGRN) images. Another DTL models work in a similar manner but just with keeping weights of the bottom layers in the feature extraction unaltered while updating weights of consecutive layers through training from scratch. Experimental results on the whole 224 SCGRN images using 5-fold cross-validation show that our model (TFeSEResNeXT101) achieving the highest average balanced accuracy (BAC) of 0.97 and thereby significantly outperforming the baseline that resulted in an average BAC of 0.86. Moreover, the simulation study demonstrated that the superiority is attributed to the distributional conformance of model weight parameters obtained with Adam optimizer when coupled with weights from a pre-trained model.

## 1. Introduction

Type 2 diabetes (T2D) is a common condition that over time when left untreated can cause damage reaching various organs, including kidney, eye, and heart, to just name a few [1, 2]. Patients with diabetes incur an overall average medical expenditure more than two times that of those without diabetes. Therefore, diabetes is considered as a burden associated with higher medical costs, and increased mortality rates [3]. Obtaining a highly accurate tool to discriminate between healthy and T2D subjects can aid in disease diagnosis, management, prevention and understanding [4]. Therefore, scientific efforts have been made contributing to detect T2D using computational methods [5-8].

Pyrros et al. [9] employed deep learning (DL)-based approach to identify T2D using chest x-ray (CXR) images pertaining to healthy and T2D patients working as follows. They employed the ResNet34 DL model incorporating typical data augmentation with the use of Adam optimizer [10]. The dataset to develop (i.e., train from scratch to induce) the ResNet34 model consisted of 271065 CXR training images in which 45961 were designated as T2D CXR images and the remaining 225104 were as CXR images for healthy control subjects. The trained model was then applied for a testing set consisting of 9943 CXR images. Results demonstrated that the DL model achieved an AUC of 0.84 when compared to an AUC of 0.79 for the baseline linear regression (LR) incorporating only clinical information data. An ensemble of both LR and ResNet34 generated an AUC of 0.85, considered as a marginal improvement in the prediction performance. These results demonstrate the feasibility of DL in screening T2D patients using CXR images. Wachinger et al. [11] presented a DL approach to predict T2D based on neck-to-knee MRI images and clinical information. The MRI images dataset consisted of 3406 MRI images in which the class distribution is uniformed (i.e., 1703 as T2D and 1703 as MRI images for healthy control subjects). The DL approach consists of convolutional layers, maxpooling layers, batch normalization layers, dropout layer and incorporation of dynamic affine feature map transform (DAFT) within convolutional layers to concatenate features obtained from clinical and MRI image data. Five-fold cross-validation was utilized to assess the performance of the whole data. Results demonstrated the superiority of CNN-DAFT achieving an AUC of 0.871, significantly outperforming CNN using only MRI images and linear regression using only clinical information.

Das [12] et al. presented a learning-based approach combining deep and machine learning for the diagnosis of T2D based on DNA sequences working as follows. First, they transformed the DNA sequence pertaining to healthy and T2D to images, provided as input to ResNet [13] and VGG19 DL models to extract features. Then, providing the extracted features along with corresponding class labels to machine learning algorithms, namely support vector machines(SVM) and KNN. Experimental results using cross-validation on the whole image dataset demonstrate the good performance of SVM when coupled with extracted features using ResNet DL model. Naveed et al. [14] employed DL to predict T2D. The dataset consisted of 19181 patient records data in which 7715 records were for diabetic patients while the remaining 11466 records were for non-diabetic patients. The dataset was divided into training and testing in which training composed of 80% of the dataset while the remaining 20% was assigned for testing. DL models included CNN, LSTM, and CNN-LSTM [15]. Experimental results demonstrated the superior performance of CNN-LSTM achieving the highest performance results when compared to other models including decision tree and SVM. Specifically, CNN-LSTM generated an accuracy of 91.6, and F1-Score of 89.2. These results demonstrate the feasibility of DL in early predicting T2D. Other AI-driven computational methods have been proposed to aid in predicting T2D [8, 16].

As inferred single-cell gene regulatory networks (SCGRNs) encode the molecular interactions pertaining to components of specific cell types and thereby can aid in characterizing cellular differentiation in healthy and disease subjects [17, 18], Turki et al. [19] presented a novel DL approach to discriminate between heathy controls and T2D based on SCGRN images working as follows. Because rapid progress in single-cell technologies has contributed to the availability of biological experiments pertaining to gene regulatory networks (GRNs), single-cell gene expression data from the ArrayExpress repository was processed with the use of bigSCale and NetBioV packages [20-22], generating 224 SCGRN images. The class distribution was distributed evenly in terms of healthy controls and T2D images. Then, utilizing RMSprop optimizer [15] with the following DL models: VGG16 [23], VGG19 [23], Xception [24], ResNet50 [13], ResNet101 [13], DenseNet121 [25], and DenseNet169 [25] to discriminate between healthy controls and T2D SCGRN images. Experimental results demonstrated the VGG19 performed better than studied DL models. However, no interpretation was provided to back the prediction performance.

Although these recently developed methods aimed to address the task of prediction T2D, these methods are still far from perfect and do not provide interpretation for practical deep transfer learning (DTL) models aiding in the explanation of the performance superiority. Therefore, this study is unique in the following aspects: (1) we present highly accurate DTL models working by combining weight parameters from pre-trained models and weights obtained with the use of Adam optimizer; (2) we provide, to the best of our knowledge, the first interpretation behind DTL models inspecting and quantifying that conformance of pre-trained model weight parameters with weight parameters obtained with the incorporation of Adam and RMSprop Optimizers. This interpretation framework can guide in the process of designing highly efficient DTL models applicable to wide range of problems; and (3) we conduct experimental study to report the prediction performance and computational running time for the task of predicting single-cell gene regulatory network images pertaining to healthy controls and T2D. Experimental results demonstrate the superiority of our DTL model, TFeSEResNeXT101, performing better than the baseline with 11% improvements. In terms of the running time, our DL models exhibited a significant reduction in training time attributed to transfer learning, which reduced the number of trainable weight parameters. In addition, simulation study unveiled the conformance of parameter weights of both transfer weights from pre-trained models with weights obtained from Adam optimizer as compared to RMSprop that was used by the baseline and resulted in inferior prediction performance, attributed to the divergence of its weight parameters from the weight parameters of pre-trained models.

### 2. Materials and Methods

### 2.1. Biological Networks

We provide an illustration in Figure 1 for the biological network images used in this study, which were downloaded from [19] and consisted of 224 SCGRN images pertaining to healthy and type 2 diabetes. The class distribution for these biological network images is balanced (i.e., 224 divided evenly into the two classes). These biological network images were produced with the help of bigSCale package to process the single-cell gene expression data and build regulatory networks, then visualizing networks via the NetBioV package. In terms of the single-cell gene expression data pertaining to healthy controls and T2D patients, it was obtained from ArrayExpress repository under accession number E-MTAB-5061 [26].

**Figure 1:**
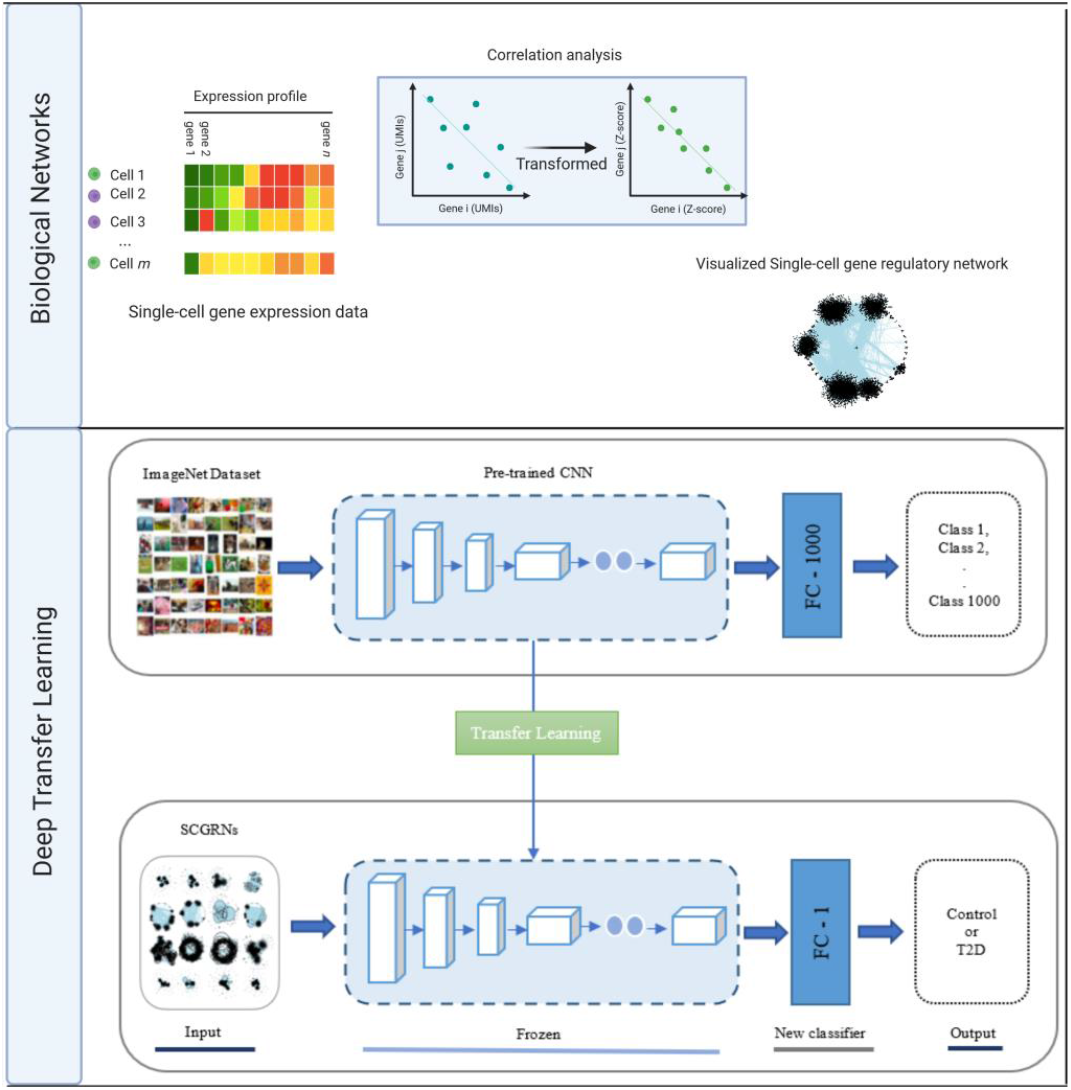
Flowchart of the deep transfer learning-based approach for the predicting T2D using SCGRNs. Biological Networks: To infer single-cell gene regulatory network (SCGRN), gene expression data are provided to bigSCale (performing clustering and differential expression analysis) changing measured correlation between genes from expression values to Z-score, followed by retaining significant correlations to guide in building a regulatory network . A visualization is performed using NetBioV. Deep Transfer Learning: Transfer learning applying feature extraction with new classifier (TFe) to distinguish between T2D and healthy control SCGRNs.

### 2.2. Deep Transfer Learning

Figure 1 demonstrates how our deep transfer learning (DTL) approach is performed. First, we adapt the following pre-trained models: VGG19, DenseNet201 [25], InceptionV3 [27], ResNet50V2 [28], ResNet101V2 [28], SEResNet152 [29], and SEResNeXt101 [29]. Each pre-trained model has a feature extraction part (i.e., series of convolutional and pooling layers) for feature extraction and a densely connected classifier for classification. Then, we keep the weights unchanged for the feature extraction part of a pre-trained model and change the densely connected classifier to deal with binary classification instead of 1000 classes. Therefore, when feeding the SCGRN image dataset, we extract features using weights of pre-trained models while training the densely connected classifier from scratch and performing prediction. We refer to models using this type of DTL computations as TFeVGG19, TFeDenseNet201, TFeInceptionV3, TFeResNet50V2, TFeResNet101V2, TFeSEResNet152, and TFeSEResNeXT101 (see Figure 1). For the other DTL computations, we keep weights of the bottom layers unchanged in the feature extraction part while performing training from scratch to change weights of top layers in feature extraction part and densely connected layers.

As in TFe-based models, we modify the densely connected classifier dealing with binary classification problem before performing the training phase. As seen in Figure 2, we refer to models employing this type of deep transfer learning as TFtVGG19, TFtDenseNet201, TFtInceptionV3, TFtResNet50V2, TFtResNet101V2, TFtSEResNet152, and TFtSEResNeXT101.

**Figure 2:**
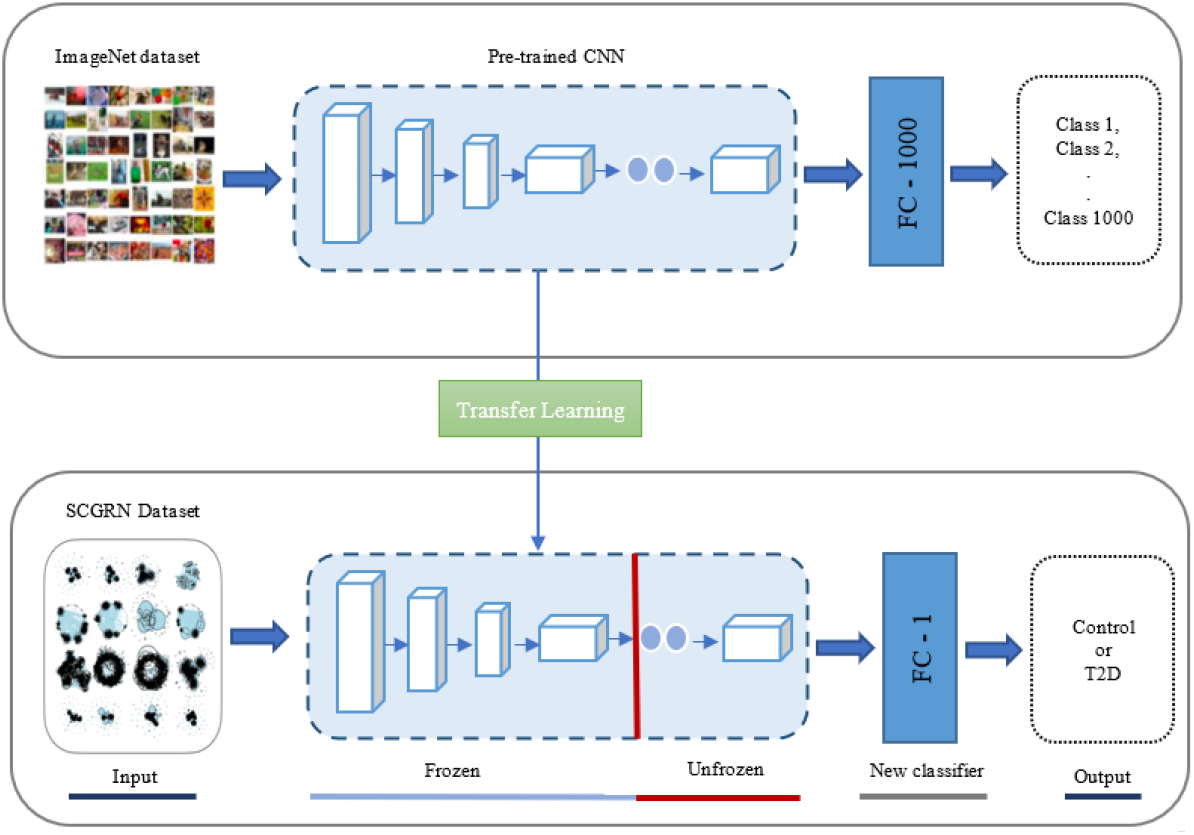
Transfer learning applying fine tuning with new classifier (TFt) to distinguish between T2D and healthy control SCGRNs.

When changing weights during training, we employed three optimizers: Adam, RMSprop, and SGD [30]. When weights are kept unchanged referring to the transfer of knowledge from pretrained models using SGD optimizer. In terms of predictions of unseen SCGRN images, predictions are mapped to healthy control subjects if the predicted values are greater than 0.5. Otherwise, predictions are mapped to T2D.

## 3. Results

### 3.2. Classification Methodology

In this study, we considered seven pre-trained models, namely VGG19, DenseNet201, InceptionV3, ResNet50V2, ResNet101V2, SEResNet152, and SEResNeXt10. Each of the pre-trained models was trained on 1.28 million images from ImageNet database to classify images into 1000 different categories. In terms of TFe-based models, we used the feature extraction part of pretrained models in which weights were kept unchanged and were used to extract feature from SCGRN images. Moreover, the densely connected classifier was trained from scratch to handle the binary class classification problem. Regarding the TFt-based models, we trained the top layers and densely connected classifier from scratch while retaining the weights of bottom layers unchanged in the feature extraction part. For both TFt-based and TFe-based models, we employed Adam optimizer when updating weights of layers. Moreover, we compared the performance of our deep transfer learning approaches using different optimizers including the baseline (i.e., RMSprop optimizer) as well as against training models from scratch. We set optimization parameters as follows: 0.00001 for the learning rate, 10 for the number of epochs, and 32 for the batch size. In terms of the loss function, we utilized categorical cross-entropy [31].

To assess the performance of studied models, we employed Balanced Accuracy (BAC), Accuracy (ACC), Precision (PRE), Recall (REC), and F1 computed as follows:

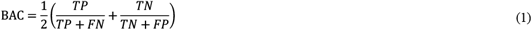

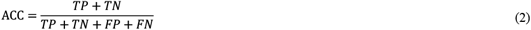

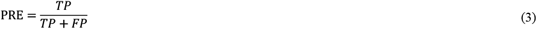

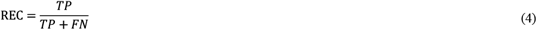

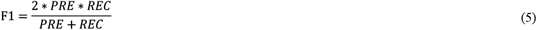

where *TN* designates true negative, corresponding to the number of T2D images that were correctly predicted as T2D. *FP* designates false positive, corresponding to the number of T2D images that were incorrectly predicted as healthy controls. *TP* designates true positive, corresponding to the number of healthy control images that were correctly predicted as healthy controls. *FN* designates false negative, corresponding to the number of healthy control images that were incorrectly predicted as T2D.

To evaluate the results on the whole SCGRN image dataset, we employed 5-fold cross-validation as follows. We partitioned the SCGRN image datasets and randomly assigned images into 5 folds. During the first run of 5-fold cross-validation, we used 4 of the folds to train our deep learning models and perform predictions to the remaining fold for testing and record the performance results. Such a process was repeated for an additional 4 runs in which performance results were recorded. Finally, we report the average performance results corresponding to the results obtained from 5-fold cross-validation.

### 3.2. Implementation Details

All experiments were run on a machine equipped with central processing unit (CPU) of Google Colab. The specifications of CPU runtime offered by Google Colab were Intel Xeon Processor with two cores with 2.30 GHz and 13 GB RAM where the installed version of Python is 3.10.11. For the analysis of models, we used R statistical software [32] to run the experiments and utilized the optimg package in R to run Adam optimizer [33]. All plots were performed using Matplotlib package in python [34].

### 3.3. Classification Results

#### 3.3.1. Training Results

In Figure 3, we illustrate the training accuracy performance results when running 5-fold crossvalidation. It can be seen that our models outperformed all other models trained from scratch. Specifically, TFeVGG19 and TFtVGG19 achieved average accuracies of 0.976 and 0.962, respectively, while VGG19 achieved an average accuracy of 0.530. TFeDenseNet201 outperformed DensNet201 via achieving an average accuracy of 0.988 while DenseNet201 performed better than TFtDenseNet201 via achieving an average accuracy of 0.982 compared to 0.946. For TFe- and TFt-based models when coupled with ResNet101V2, SEResNet152 and SEResNetXT101, they outperformed their counterparts when not applying deep transfer learning (DTL) models. These superior performance results are attributed to the learned representation using transfer learning.

**Figure 3:**
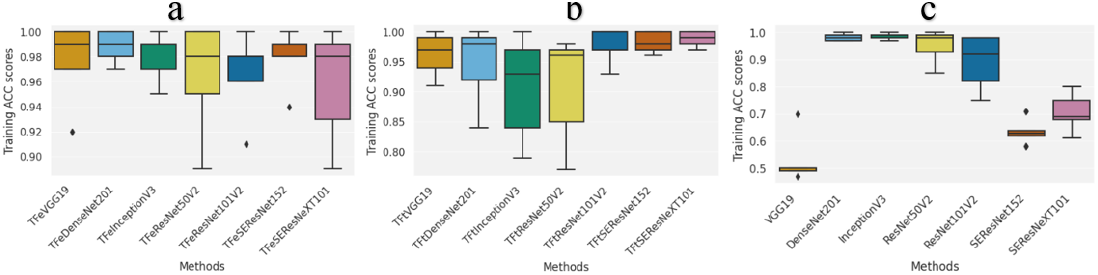
The boxplots presenting the average 5-fold cross-validation results using the ACC measure for the training folds. (a) Deep transfer learning models using feature extraction (referred with the prefix TFe). (b) Deep transfer learning models using fine tuning (referred with the prefix TFt). (c) Deep learning models trained from scratch. ACC is accuracy.

#### 3.3.2. Testing Results

Figures 4 and 5 report the generalization (i.e., test) accuracy performance results and combined confusion matrices, respectively, when 5-fold cross-validation is utilized. TFeSeResNeXT101 achieved the highest average accuracy of 0.968.

**Figure 4:**
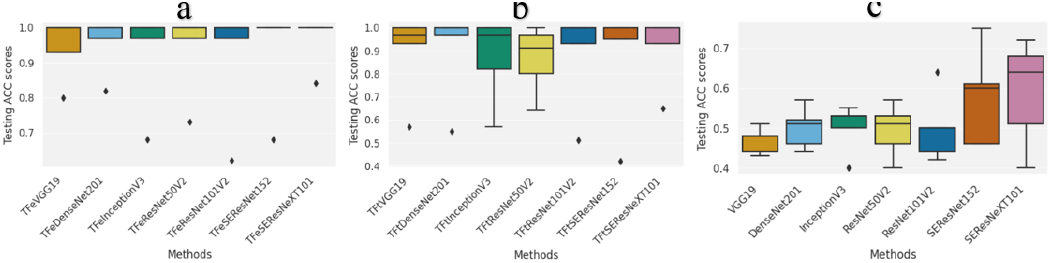
The boxplots presenting the average 5-fold cross-validation results using the ACC measure for the testing folds. (a) Deep transfer learning models using feature extraction (referred with the prefix TFe). (b) Deep transfer learning models using fine tuning (referred with the prefix TFt). (c) Deep learning models trained from scratch. Acc is accuracy.

**Figure 5:**
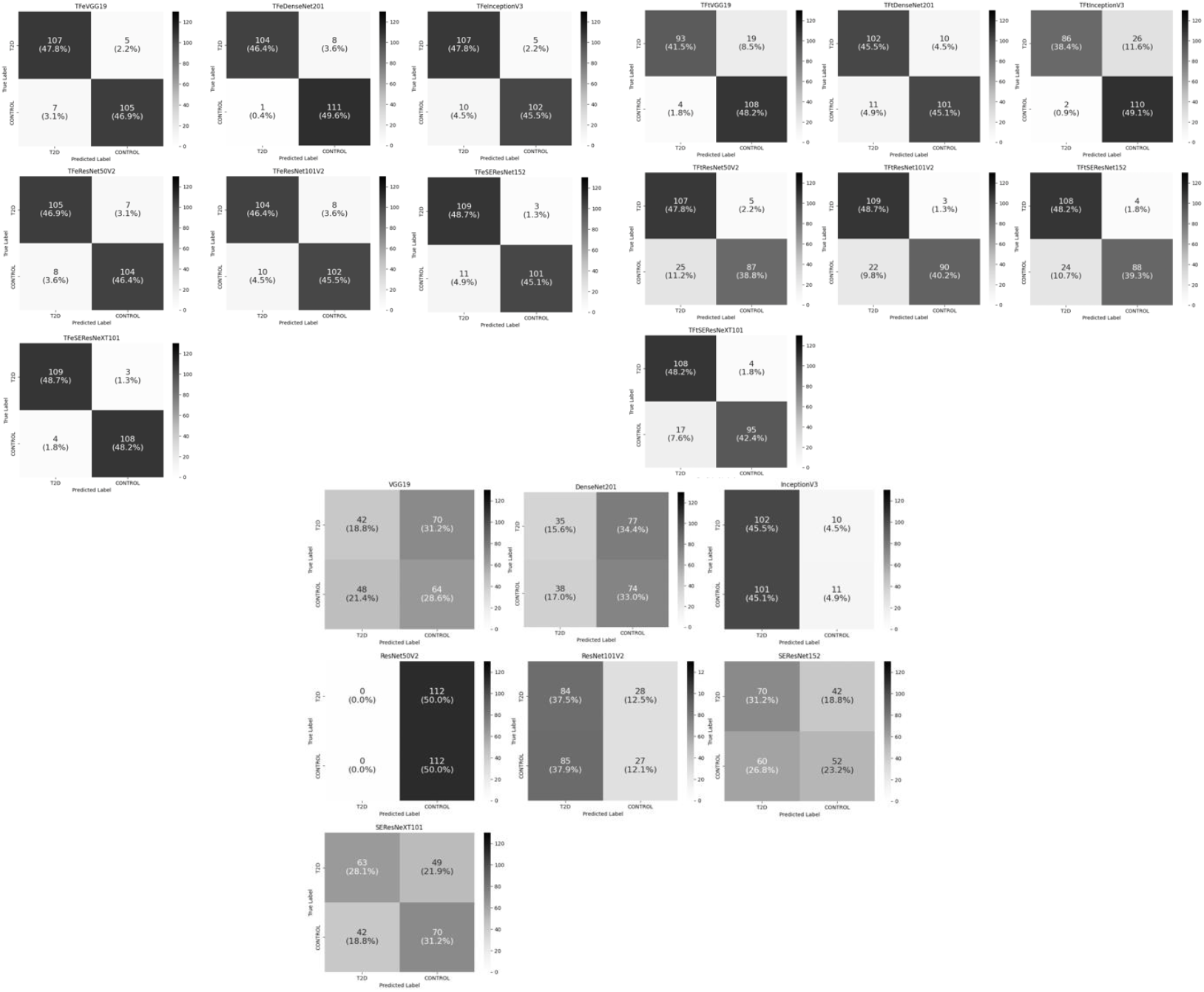
Combined confusion matrices for all methods during the running of 5-fold cross-validation.

The second-best model is TFeDenseNet201, achieving an average accuracy of 0.958, followed by TFeVGG19, TFeResNet50V2, TFeSEResNet152, TFeInceptionV3, and TFeResNet101V2 (generating average accuracies of 0.946, 0.940, 0.936, 0.930, and 0.918, respectively). TFt-based models also outperformed all models trained from scratch (see Figure 4(b and c)). Particularly, TFt-based models generated average accuracies lower and upper bounded by 0.864 and 0.916, respectively, while models trained from scratch were lower and upper bounded by average accuracies of 0.468 and 0.590. These results demonstrate the superior performance of models employing our DTL computations.

In terms of reporting testing performance results using different metrics, our model TFeSeResneXT101 outperforms all other models (see Table 1) via achieving an average BAC of 0.97, average PRE of 0.97 (tie with our model TFeSEResNet152), and average F1 of 0.97. Moreover, TFeVGG19 and TFtVGG19 perform better than VGG19. Similarly, TFeDenseNet201, TFeInceptionV3, TFeResNet50V2, TFeResNet101V2, and TFeSEResNet152 performed better than DenseNet201, InceptionV3, ResNet50V2, ResNet101V2, and SEResNet152, respectively. Th same holds true for TFt-based models outperforming their counterparts (i.e., VGG19, Dense-Net201, InceptionV3, ResNet50V2, ResNet101V2, SEResNet152, and SEResNetXT101)

**Table 1:** Reported average performance results during the running of 5-fold cross-validation on testing using studied models. BAC is balanced accuracy. PRE is precision. REC is recall. The best overall result is underlined and is shown in bold. The method outperforming its counterparts is just underlined.

| Model | Optimizer | BAC | PRE | REC | F1 |
| --- | --- | --- | --- | --- | --- |
| TFeVGG19 | Adam | <u>0.94</u> | <u>0.95</u> | 0.94 | <u>0.94</u> |
| TFtVGG19 | Adam | 0.90 | 0.82 | **<u>0.97</u>** | 0.89 |
| VGG19 | Adam | 0.50 | 0.37 | 0.46 | 0.41 |
| TFeDenseNet201 | Adam | <u>0.96</u> | <u>0.95</u> | <u>0.95</u> | <u>0.96</u> |
| TFtDenseNet201 | Adam | 0.90 | 0.91 | 0.90 | 0.90 |
| DenseNet201 | Adam | 0.50 | 0.20 | 0.51 | 0.29 |
| TFeInceptionV3 | Adam | <u>0.93</u> | <u>0.95</u> | 0.91 | <u>0.92</u> |
| TFtInceptionV3 | Adam | 0.87 | 0.76 | <u>0.92</u> | 0.83 |
| InceptionV3 | Adam | 0.50 | 0.91 | 0.50 | 0.64 |
| TFeResNet50V2 | Adam | <u>0.93</u> | 0.93 | <u>0.92</u> | <u>0.93</u> |
| TFtResNet50V2 | Adam | 0.87 | <u>0.95</u> | 0.81 | 0.87 |
| ResNet50V2 | Adam | 0.50 | - | - | - |
| TFeResNet101V2 | Adam | <u>0.92</u> | <u>0.92</u> | <u>0.91</u> | <u>0.92</u> |
| TFtResNet101V2 | Adam | 0.89 | 0.90 | 0.86 | 0.88 |
| ResNet101V2 | Adam | 0.52 | 0.75 | 0.49 | 0.59 |
| TFeSEResNet152 | Adam | <u>0.94</u> | **<u>0.97</u>** | <u>0.90</u> | <u>0.93</u> |
| TFtSEResNet152 | Adam | 0.88 | 0.96 | 0.81 | 0.88 |
| SEResNet152 | Adam | 0.55 | 0.67 | 0.50 | 0.57 |
| TFeSEResNeXT101 | Adam | **<u>0.97</u>** | **<u>0.97</u>** | <u>0.96</u> | **<u>0.97</u>** |
| TFtSEResNeXT101 | Adam | 0.90 | 0.96 | 0.90 | 0.91 |
| SEResNeXT101 | Adam | 0.60 | 0.56 | 0.60 | 0.58 |

Table 2 reports our best DTL models with different optimizers. It can be shown that TFeSeResNeXT101 and TFtSeResNeXT101 generate the highest performance results when coupled with Adam optimizer method. Specifically, TFeSEResNeXT101 with Adam optimizer generates the highest average BAC (and F1) of 0.97 (and 0.97). TFtSEResNeXT101 with Adam optimizer archives the highest average BAC of 0.91, highest average F1 of 0.90 (tie with SGD optimizer). When TFeSEResNeXT101 and TFtSEResNeXT101 are coupled with SGD optimizer, they generate inferior performance results.

**Table 2:** Performance comparison of our best deep transfer learning model under different optimizers during the 5-fold cross-validation. BAC is balanced accuracy. Best performance result is shown in bold.

| Model | Optimizer | BAC | F1 |
| --- | --- | --- | --- |
| TFeSEResNeXT101 | Adam | **0.97** | **0.97** |
|  | RMSprop | 0.92 | 0.92 |
|  | SGD | 0.77 | 0.77 |
| TFtSEResNeXT101 | Adam | **0.91** | **0.90** |
|  | RMSprop | 0.90 | 0.89 |
|  | SGD | 0.90 | **0.90** |

In Table 3, we compare our model TFeVGG19 with Adam optimizer against the best performing baseline TFeVGG19 with RMSprop optimizer, named VGG19 in [19]. It is evident that our model TFeVGG19 with Adam optimizer achieves the highest average BAC of 0.94 while the baseline obtained an average BAC of 0.86. Moreover, when F1 performance measure is considered, TFeVGG19 with Adam optimizer attains the highest average F1 of 0.94 while the baseline achieved an average F1 of 0.88. The same holds true for TFtVGG19, which achieved the highest average BAC of 0.91, highest average F1 of 0.90.

**Table 3:**
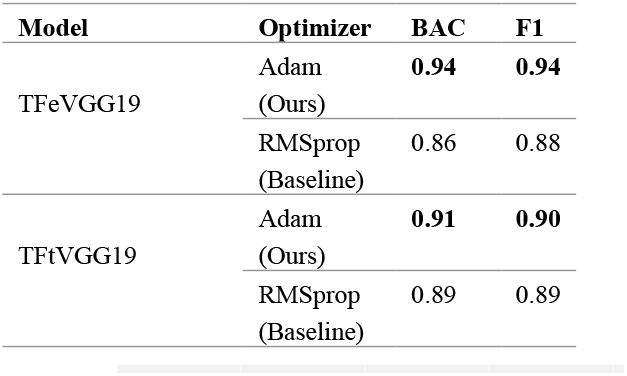
Performance comparison of our deep transfer learning model against recent baseline methods when 5-fold cross-validation is employed. BAC is balanced accuracy. Best performance result is shown in bold.

In Figure 6, we report the running time in seconds for the process of running 5-fold crossvalidation when utilizing our best model (TFeSEResNeXT101) and TFtSEResNeXT101 compared to their peer SeResneXT101. Our model TFeSEResNeXT101 is 208.45x faster than SEResNeXT101. Also, our model TFtSEResNeXT101 is 3.82x faster than SEResNeXT101. Moreover, TFeVGG19 and TFtVGG19 are 802.67x and 2.53x, respectively, faster than VGG19. These results demonstrate the computational efficiency of the DTL models, in addition to the highly achieved performance results.

**Figure 6:**
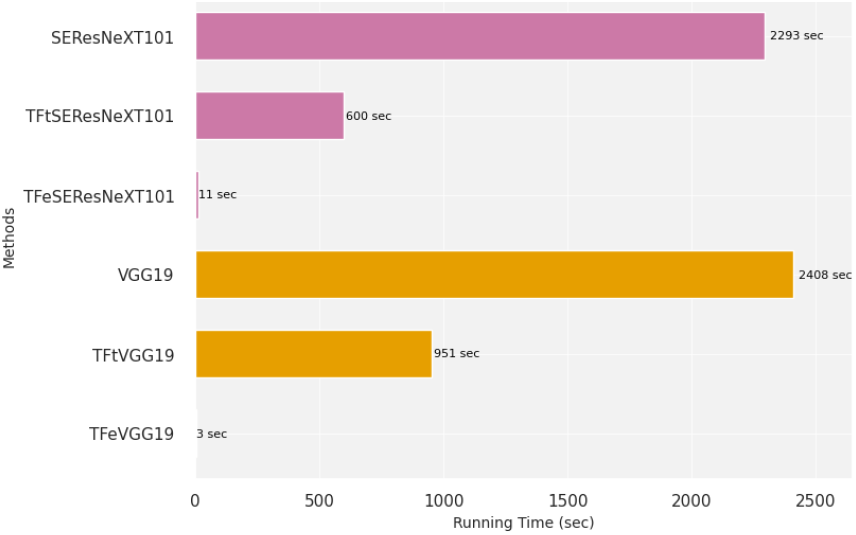
Running time comparisons in seconds for selected models when running 5-fold cross-validation.

### 3.4. Models Introspection

#### 3.4.1. Stochastic Gradient Descent (SGD)

To minimize the objective function *Q*(*θ*_0_, *θ*_1_) for parameters *θ*_0_ an*d θ*_1_ of model *H*(*x*_*i*_), we employ gradient descent optimization algorithms to find *θ*_0_ an*d θ*_1_ minimizing the objective function. The optimization problem can be formulated as follows:

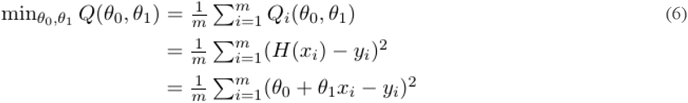

We utilize SGD, RMSprop, and Adam optimization algorithm to minimize the objective function and estimate the model parameters. For SGD, we initialize the parameters *θ*_0_ an*d θ*_1_ according to the uniform distribution *U*(0,1) and setting the learning rate η = 0.001, maximum number of iterations to 3000. Then, in each time, shuffling the data of *m* examples followed by looping *m* times over the following to update model parameters After the end of looping, the algorithm stops if the maximum number of iterations is reached or ‖∇*Q*(*θ*_0_, *θ*_1_)‖ ≤ 0.001:

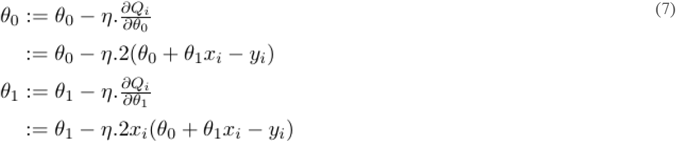

#### 3.4.2. Root Mean Square Propagation (RMSprop)

For RMSprop, we initialize model parameters according to uniform distribution *U*(0,1), set the learning rate η = 0.001, maximum number of iterations to 3000, *β* = 0.9, *∈* = 10^−6^, Batch-Size = 16, referred in the following as |*S*|. Then, looping to update model parameters according to each selected batch. After ending of looping over all selected batches, the algorithm terminates when ‖∇*Q*(*θ*_0_, *θ*_1_)‖ ≤ 0.001 or the maximum number of iterations is reached:

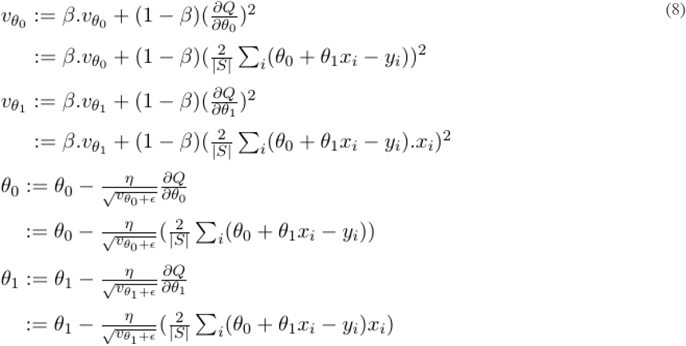

#### 3.4.3. Adaptive Moment Estimation (Adam)

In terms of Adam, 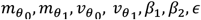, and η are initialized as in [35]. Then, looping to update model parameters according to all selected batches of examples. After ending of looping over all selected batches, the algorithm stops if the maximum number of iterations (i.e., 3000) is reached or ‖*▽Q*(*θ*_0_, *θ*_1_)‖ ≤ 0.001:

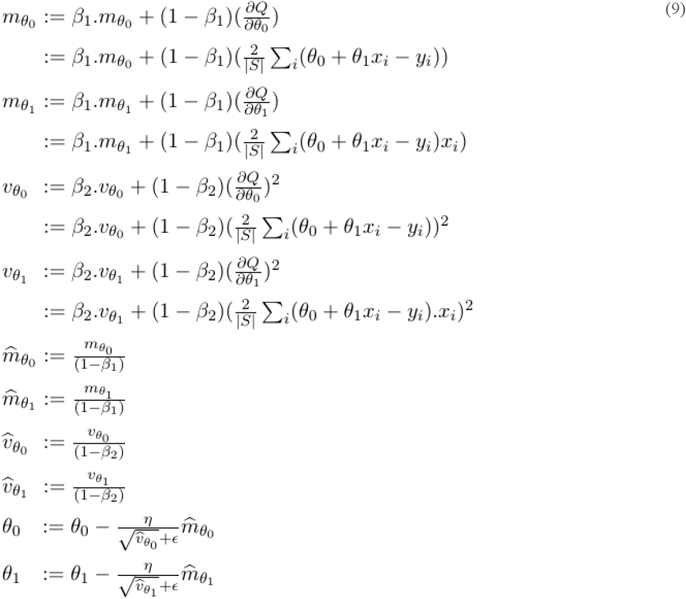

#### 3.4.4. Simulated Data

To demonstrate the efficiency of the proposed deep transfer learning (DTL) models incorporation mixed parameters derived from both SGD and Adam, we conducted simulation studies to explain the superiority behind the proposed models as well as imitate the numerical behavior. Particularly, we consider the following four predictive models:

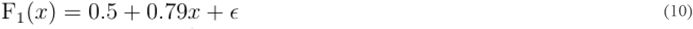

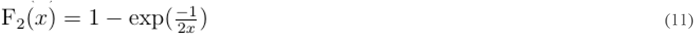

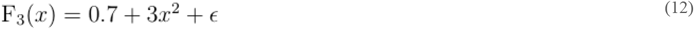

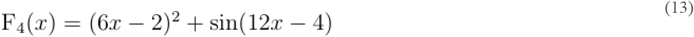

where *X*∼*U*(0,1) and *∈*∼*N*(0,0.2) in which *U*() and *N*() are uniform and normal distributions, respectively. For F_1_, when we have *X* and *Y*=F_1_(*X*), we perform the following steps. Let *H*_*SGD*_(*x*_*i*_) = *θ*_0_ *+ θ*_1_*x*_*i*_ (for *i* =1..*m*) be the model in which we want to estimate parameters using (*X,Y*) data from F_1_ coupled with Equation 7. Similarly, let *H*_*RMSprop*_(*x*_*i*_) and H_*A*d*am*_(*x*_*i*_) be models in which we want to estimate their parameters using Equations 8 and 9, respectively, coupled with (*X,Y*) data from F_1_. Then, we provide each *x*_*i*_ *∈* X to perform predictions corresponding to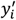 . In Figure 7, we report 2D plots for *X* and predicted 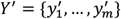 via each model using data generated according to Equation 10, where SGD refers to plotting (*x*_*i*_, H_*SGD*_(*x*_*i*_)) while Adam and RMSprop refers to plotting results obtained via (*x*_*i*_, H_*A*d*am*_ (*x*_*i*_)) and (*x*_*i*_, H_*RMSprop*_(*x*_*i*_)), respectively, and *i*=1..*m*. We then repeat this process for an additional 8 runs. Therefore, we have 9 runs in total It can be seen from Figure 7 that model induced via RMSprop has more distributional differences compared to those obtained via SGD and Adam. To quantify distributional differences between SGD and Adam against SGD and RMSprop, we perform the following computations:

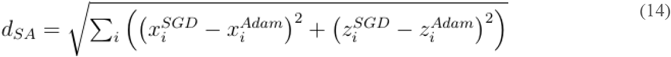

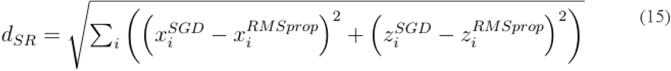

where *d*_SA_ measuring the distance between data associated with SGD and Adam. Similarly, *d*_SR_ measures the distance between data associated with SGD and RMSprop. The lower the distance value, the less the distribution difference is. Figure 11(a) plots *d*_SA_ and *d*_SR_ for the 9 runs. It can be seen that *H*_*A*d*am*_ (*x*_*i*_) is closer to *H*_*SGD*_(*x*_*i*_) than *H*_*RMSprop*_(*x*_*i*_) to *H*_*SGD*_(*x*_*i*_) in most runs. Moreover, the distributional differences are statistically significant (*P*-value = 7.28 × 10^−14^ from *t*-test).

**Figure 7:**
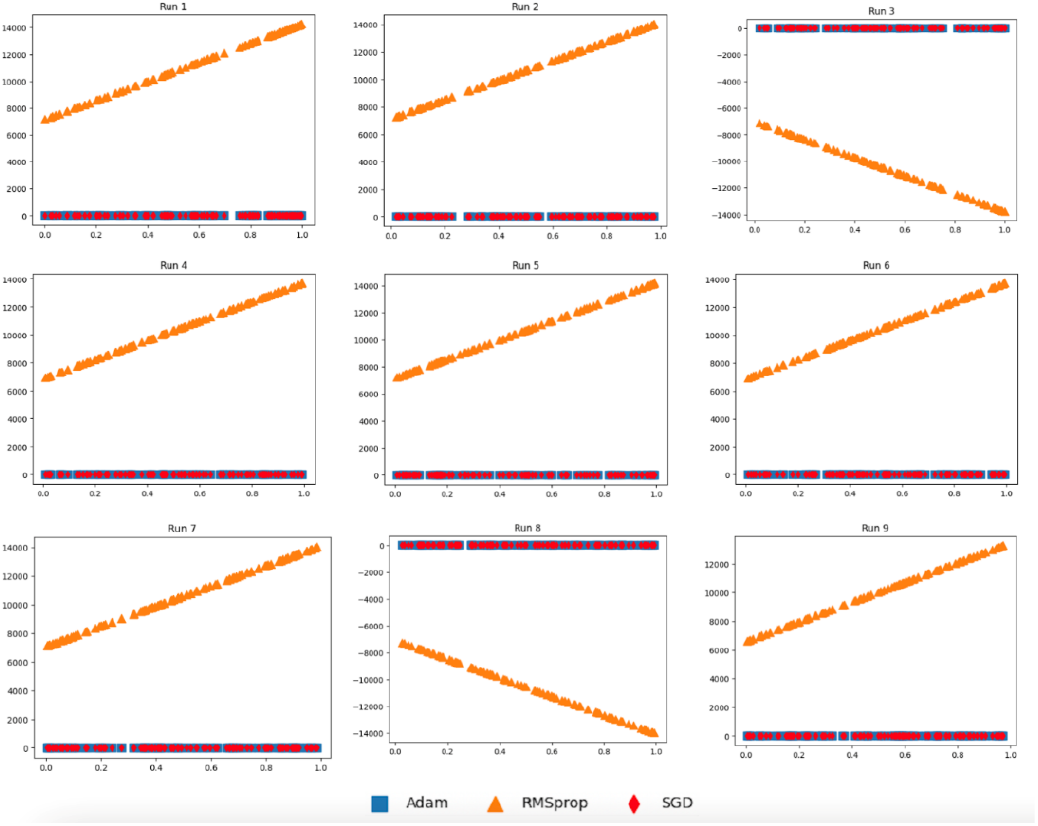
Plots for the three models as (*x*_*i*_, H_*SGD*_(*x*_*i*_)), (*x*_*i*_, H_*A*d*am*_ (*x*_*i*_)), and (*x*_*i*_, H_*RMSprop*_(*x*_*i*_)) for *i=*1..*m* according to *x*_*i*_ generated using F_1_.

These results demonstrate conformance of the weight parameters of models utilizing Adam and SGD optimizers. Figure 8 reports the 2D plots of three induced models as (*x*_*i*_, H_*SGD*_(*x*_*i*_)), (*x*_*i*_, H_*A*d*am*_ (*x*_*i*_)), *and* (*x*_*i*_, H_*RMSprop*_(*x*_*i*_)) for *i*=1..*m* using data generated according to Equation 11 (i.e., F_2_) [36]. It can be clearly seen that the data distributional difference of results via SGD is closer to that of Adam when compared to results obtained with the help of RMSprop. In Figure 11(b), we quantify distributional differences using Equations 14 and 15. It can be shown that Adam is closer to SGD as shown from AdamSGD when compared to that of RMSprop to SGD (i.e., RMSpropSGD) over the 9 runs. The quantification of AdamSGD is attributed to *d*_SA_ while RMSpropSGD is attributed to *d*_SR._ In addition, the distributional differences between AdamSGD and RMSpropSGD are statistically significant (*P-*value = 7.01 × 10^−7^ from *t*-test).

**Figure 8:**
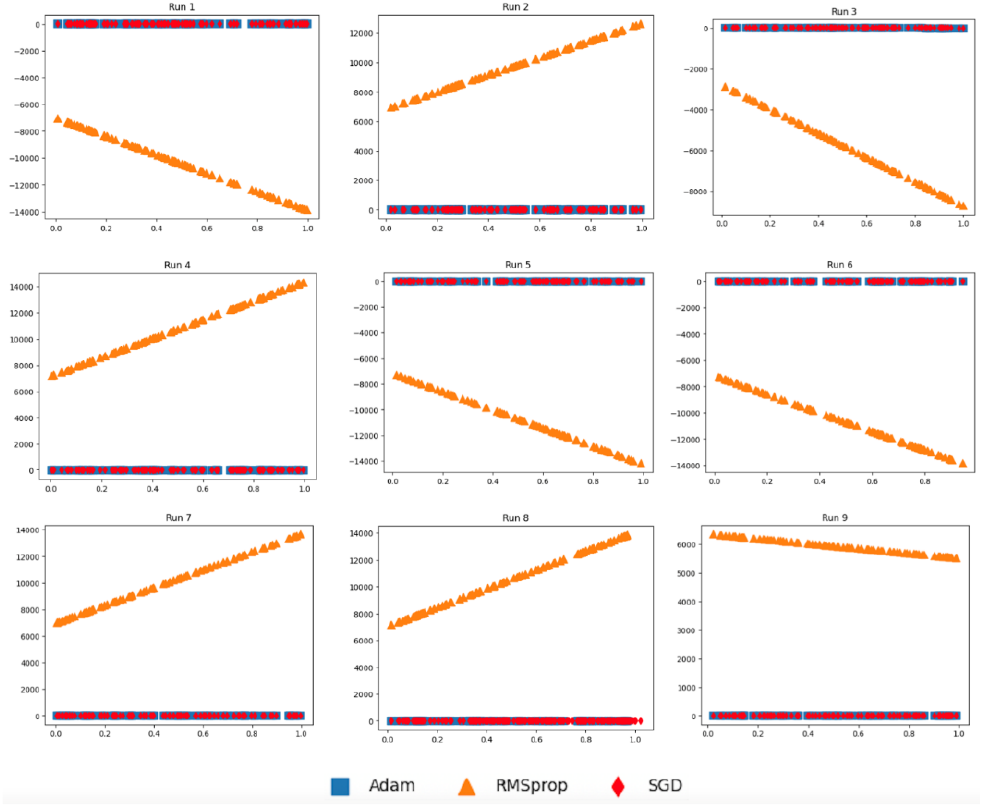
Plots for the three models as (*x*_*i*_, *H*_*SGD*_(*x*_*i*_)), (*x*_*i*_, *H*_*Adam*_(*x*_*i*_)), and (*x*_*i*_, *H*_*RMSprop*_(*x*_*i*_)) for *i*=1..*m* according to *x*_*i*_ generated using F_2_.

Figures 9 and 10 report 2D plots of (*x*_*i*_, H_*SGD*_(*x*_*i*_)), (*x*_*i*_, H_*A*d*am*_ (*x*_*i*_)), *and* (*x*_*i*_, H_*RMSprop*_(*x*_*i*_)) for *i*=1..*m* using generated data of Equations 12 (F_3_) and 13 (F_4_) [37], where models were induced with SGD, Adam, and RMSprop optimizers. It can be seen from the alignment of Adam with SGD that Adam has a closer data representation to SGD compared to RMSprop to SGD. When quantifying the data distributional differences in Figure 11 (c and d), it can be clearly shown that the distributional differences of SGD and Adam (referred to AdamSGD) are closer than SGD to RMSprop over the 9 runs.

**Figure 9:**
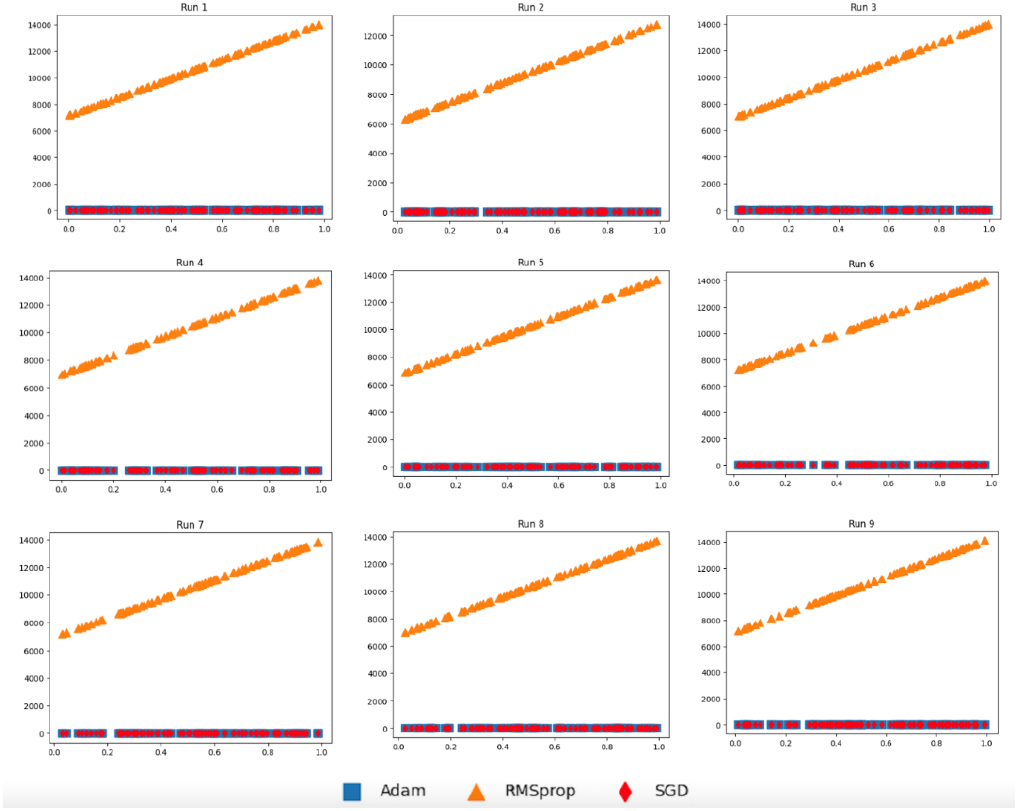
Plots for the three models as (*x*_*i*_, *H*_*SGD*_(*x*_*i*_)), (*x*_*i*_, *H*_*Adam*_(*x*_*i*_)), and (*x*_*i*_, *H*_*RMSprop*_(*x*_*i*_)) for *i*=1..*m* according to *x*_*i*_ generated using F_3_.

**Figure 10:**
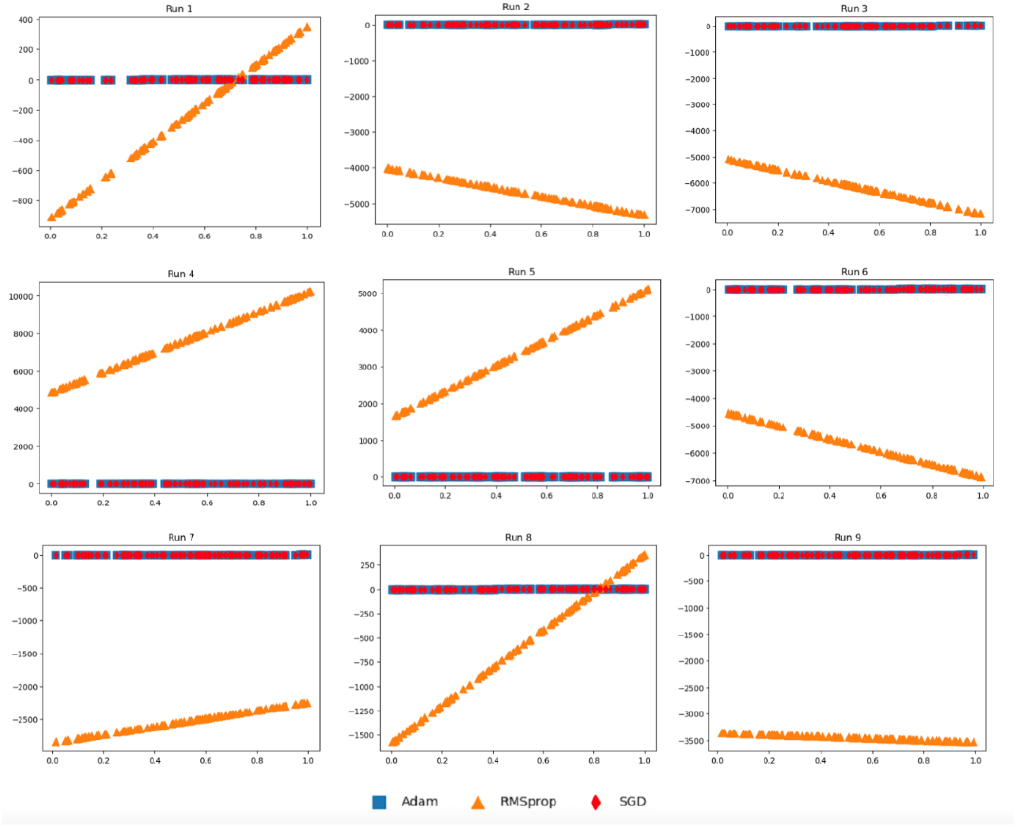
Plots for the three models as (*x*_*i*_, H_*SGD*_(*x*_*i*_)), (*x*_*i*_, H_*Adam*_(*x*_*i*_)), and (*x*_*i*_, H_*RMSprop*_(*x*_*i*_)) for *i*=1..*m* according to *x*_*i*_ generated using F_4_.

These quantified results for AdamSGD and RMSprop are attributed to *d*_SA_ and *d*_SR_, respectively. Additionally, the distributional differences of between AdamSGD and RMSprop were statistically significant (*P-*value = 3.48 × 10^−12^ from *t*-test when F3 is used while *P-*value = 1.49 × 10^−3^ from *t*-test when F4 is used). These results demonstrate the stable performance when SGD is coupled with Adam.

## 4. Discussion

Our deep transfer learning (DTL) models work as follows. In the TFe-based models, the convolutional base (also called the feature extraction part) in the pre-trained model is left unchanged while the densely connected classifier is modified to deal with the binary class classification at hand. Therefore, we applied the features extraction part of pre-trained models to the SCGRN images to extract features followed by a flattening step to train densely connected classifier from scratch. It can be noted that only weights of densely connected classifier are changed according to Adam optimizer while we transferred knowledge (i.e., weights) of the feature extraction part from pre-trained models. In terms of the TFt-based models, we keep weights of the bottom layers in the feature extraction part of pre-trained models unchanged while modifying weights in the proceeding layers including the densely connected classifier according to the Adam optimizer. Moreover, the densely connected classifier was altered to deal with the binary class classification problem pertaining to distinguishing between healthy controls and T2D SCGRN images. It can be seen that updating model weight parameters is done through the training with the use of Adam optimizer.

When conducting experimental study to assess deep transfer learning models, we used different optimizers, including SGD, RMSprop, and Adam. For SGD optimizer used in our DTL models, the model weight parameters were different than DTL models coupled with RMSprop and Adam optimizers. Therefore, we induced three sets of different models attributed to the three optimizers. Experimental results demonstrate the superiority of DTL models utilizing SGD and Adam optimizers when compared to that using SGD and RMSprop optimizers. We reported training loss for epochs pertaining to TFe-based and TFt-based models when running 5-fold cross validation in Supplementary Figure S1. For each DTL model, the number of layers including frozen and unfrozen layers is reported in Supplementary Table S1.

In our study, mitigation of overfitting is attributed to (1) transfer learning in which many layers in DTL models are freezed and thereby reducing the number of trainable parameters; and (2) applying the dropout to the last fully-connected layer in which we set the dropout rate to 0.5 [38]. It is worth mentioning that we assessed the performance of other deep learning (DL) models such as ConvNeXtTiny and ConvNeXtLarge [39]. Although ConvNeXtLarge outperformed ConvNeXtTiny, they didn’t exhibit superior performance when compared to TFeSEResNeXT101. Therefore, we include their performance results in Supplementary Tables S2 and S3 (and Supplementary Figure S2). In terms of the running time, TFeSEResNeXT101was 1.54x faster than TFeConvNeXtLarge. We report running time for ConvNeXtTiny-based and ConvNeXtLargebased models in Supplementary Figure S3.

In our DTL models, we have transferred weights from pre-trained models coupled with weights obtained with the help of Adam optimizer. If weight parameters obtained using two optimizers are close, then the two models almost behave the same. On the other hand, when the weight parameters resulted from two optimizers are not close, then the two models behave differently. To mimic the real scenario and investigate the effects of coupling different model weight parameters, we performed a simulated study. In Figures 7-11, we showed that a model induced with the help of SGD optimizer is closer to a model induced with Adam optimizer when compared to a model induced with the help of RMSprop optimizer. It can be evident from visualized results in our study that SGD and Adam had less distributional differences than that of SGD and RMSprop. That resembles the case of having two related datasets for SGD and Adam against unrelated datasets for SGD and RMSprop. As a result, inferior performance results for models utilizing RMSprop are attributed to the high distributional differences in model weight parameters.

**Figure 11:**
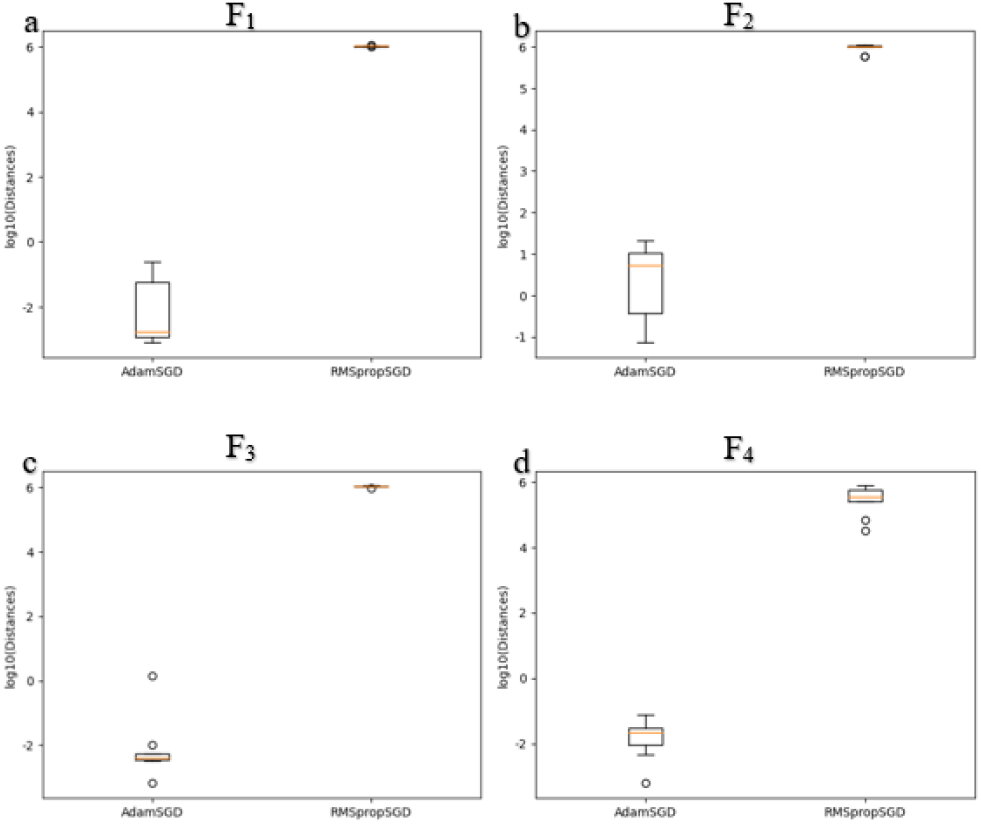
Boxplots of the four studied models, F_1_-F_4_, showing the distance distribution over nine runs for AdamSGD and RMSpropSGD. a) results for F_1_. B) results for F_2_. c) results for F_3_. d) results for F_4_.

It is worth noting that our DTL models keep weights of many layers unchanged. Therefore, when we trained our models, we had fewer number of updated weights compared to updated weights in models trained from scratch. It can be seen from Figure 6 that our DTL models are fast and can be adopted into mobile applications. It can be noticed from Tables 2 and 3 that leveraging source task knowledge contributed to improved prediction performance when coupled with updated weight parameters in the target task using Adam optimizer. On the other hand, the transferred knowledge from the source task contributed to degraded performance when coupled with updated weight parameters in the target task using RMSprop and SGD optimizers. Also, when we assessed additional DL models such as ConvNeXtLarge and ConvNeXtTiny, the knowledge transfer contributed to maintain the same performance behavior in which leveraging source domain knowledge when coupled with updated weights in the target task remained to be the best (see Supplementary Tables S2 and S3).

## 5. Conclusions and Future Work

In this paper, we present and analyze deep transfer learning (DTL) models for the task of classifying 224 SCGRN images pertaining to healthy controls and T2D patients. First, we utilized seven pre-trained models (including SEResNet152 and SEResNeXt101) already trained on more than million images from the ImageNet dataset. Then, we left weights in the convolutional base (i.e., feature extraction part) unchanged and thereby transferring knowledge from pre-trained models while modifying the densely connected classifier with the use of Adam optimizer to discriminate heathy and T2D SCGRN images. Another presented DTL models work as follows. We kept weights of bottom layers in the feature extraction part of pre-trained model unchanged while modifying consequent layers including the densely connected classifier with the use of Adam optimizer. Experimental results on the whole 224 SCGRN image dataset using 5-fold cross-validation demonstrate the superiority of TFeSEResNeXT101, achieving the highest average BAC of 0.97 and therefore significantly surpassing the performance of the baseline resulted in an average BAC of 0.86. Furthermore, our simulation study showed that the highly accurate performance in our models is attributed to the distributional conformance of weights obtained with the use of Adam optimizer when coupled with weights of pre-trained models.

Future work includes (1) adopting our computational framework to analyze DTL models with different network topologies and thereby identifying the best practice for DTL; (2) incorporating multi-omics datasets with images to improve the prediction performance using DTL models; (3) developing a boosting mechanism to improve the performance of DTL models in different biological problems [40, 41]; (4) incorporating feature representation obtained via our DTL models with machine learning algorithms for the task of inferring SCGRNs; and (5) utilizing our framework to speed up the learning process, e.g., TFeVGG19 was 802.67x faster than VGG19, trained from scratch.

## Author Contributions

T.T. conceived and designed the study. S.A. performed the deep learning experiments and the visualization of results. T.T. performed the analysis. T.T. and S.A. wrote the manuscript. T.T. supervised the study. All authors have read and agreed to the revised version of the manuscript.

## Data Availability

The dataset analyzed during the current study is available in the dataset folder within supplementary material at https://www.biorxiv.org/content/10.1101/2020.08.30.273839v1.supplementary-material. The single-cell gene expression data is available in the ArrayExpress repository under accession number E-MTAB-5061 (https://www.ebi.ac.uk/biostudies/arrayexpress/studies/E-MTAB-5061).

