## Supplementary material for "A novel interpretable deep transfer learning combining diverse learnable parameters for improved T2D prediction based on single-cell gene regulatory networks": Supplementary Figures.docx

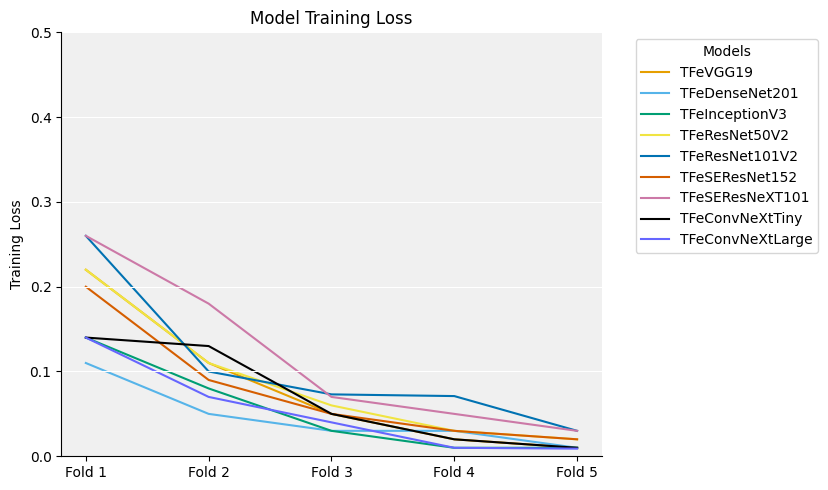

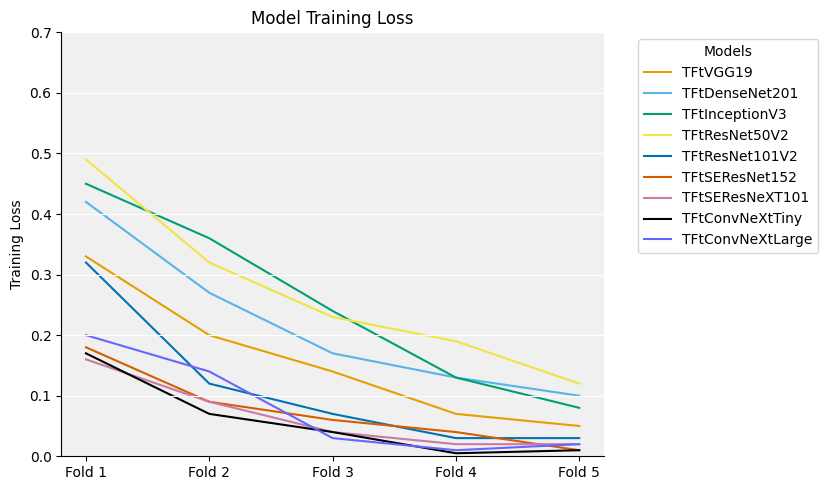

**Figure S1**: Reported training loss for epochs pertaining to TFe-based and TFt-based models when running 5-fold cross validation. TFe is transfer learning applying fine extraction with new classifier. TFt is transfer learning applying fine tuning with new classifier.

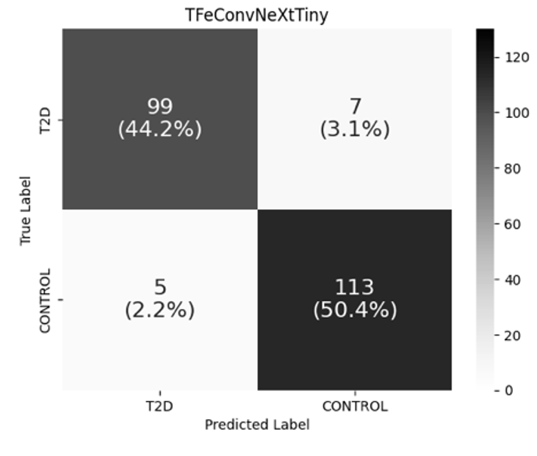

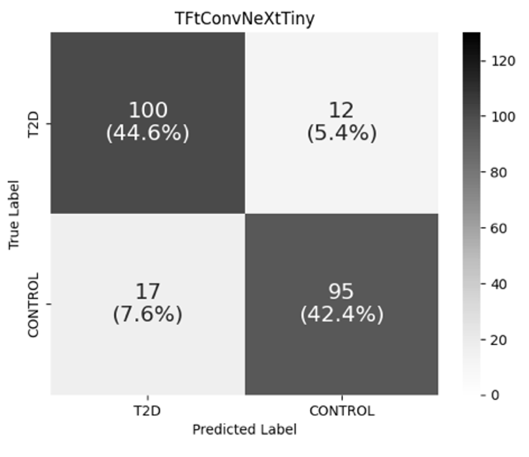

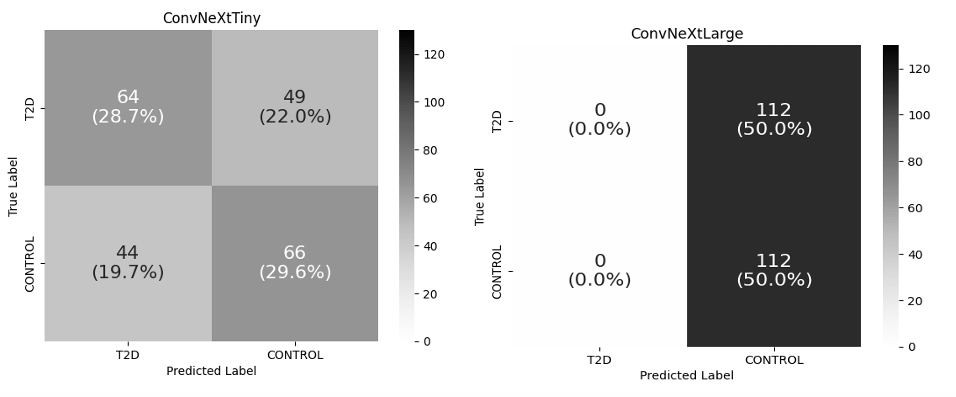

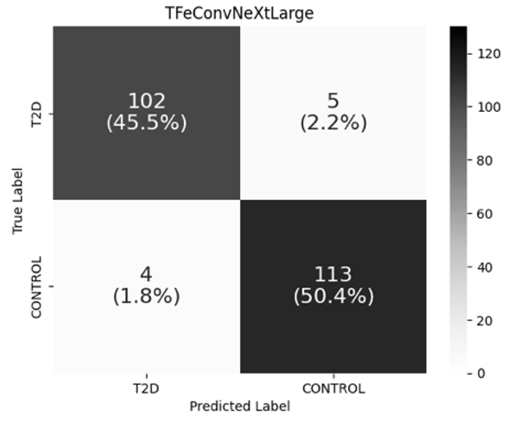

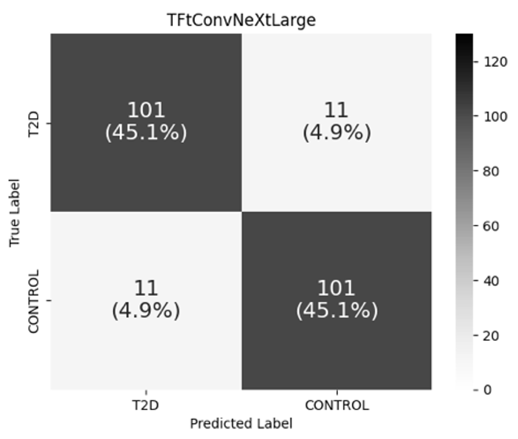

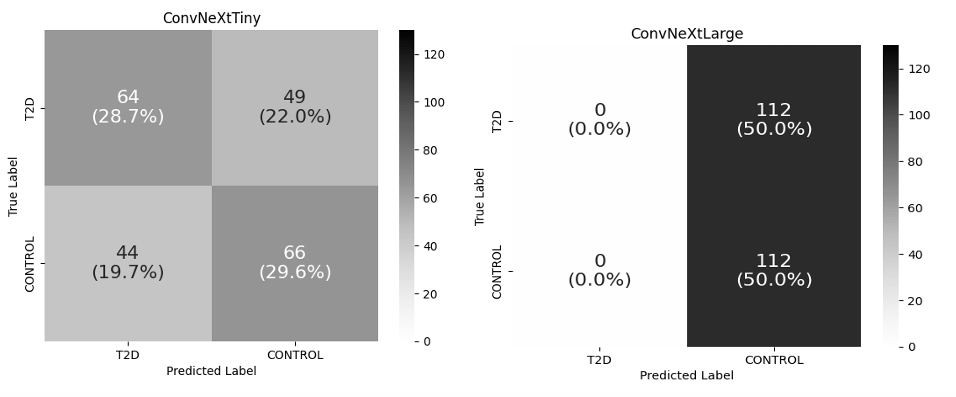

**Figure S2**: Combined confusion matrices for ConvNeXtTiny-based and ConvNeXtLarge-based models during the running of 5-fold cross-validation.

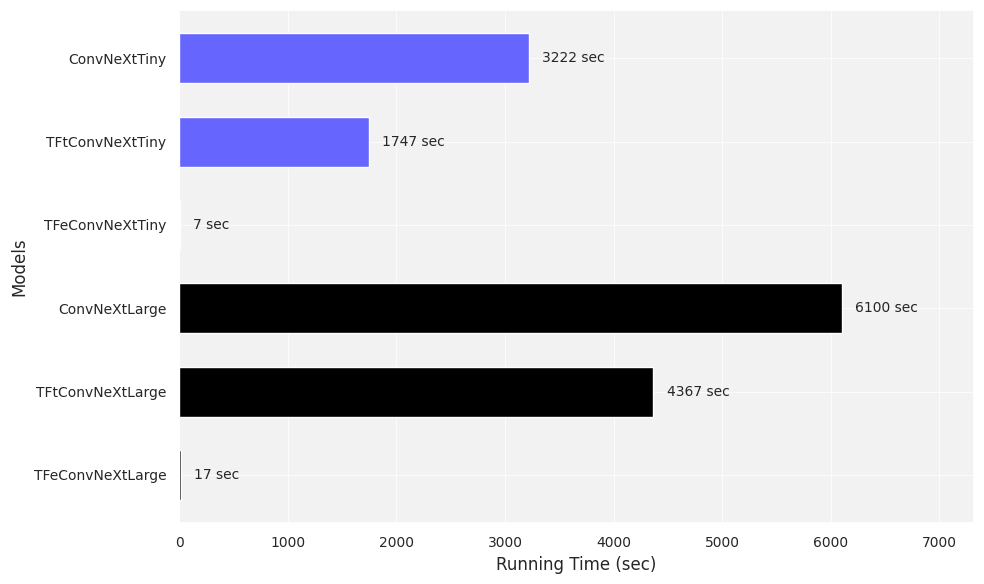

**Figure S3*:*** Running time comparisons in seconds for ConvNeXtTiny-based and ConvNeXtLarge-based models when running 5-fold cross-validation.
