## Supplementary material for "A novel interpretable deep transfer learning combining diverse learnable parameters for improved T2D prediction based on single-cell gene regulatory networks": Supplementary Tables.docx

| Model | Number of layers | Number of Frozen layers | Number of Unfrozen layers |
| --- | --- | --- | --- |
| TFtVGG19 | 22 | 14 | 8 |
| TFtDenseNet201 | 707 | 500 | 207 |
| TFtInceptionV3 | 311 | 280 | 31 |
| TFtResNet50V2 | 190 | 170 | 20 |
| TFtResNet101V2 | 377 | 300 | 77 |
| TFtSEResNet152 | 915 | 800 | 115 |
| TFtSEResNeXT101 | 2721 | 2300 | 421 |
| TFtConvNeXtTiny | 151 | 130 | 21 |
| TFtConvNeXtLarge | 295 | 255 | 40 |

**Table S1**: Number of layers including frozen and unfrozen layers during the development of TFt-based models.

**Table S2**: Performance comparison of deep transfer learning models employing ConvNextLarge under different optimizers during the 5-fold cross-validation. BAC is balanced accuracy. Best performance result is shown in bold.

| Model | Optimizer | BAC | F1 |
| --- | --- | --- | --- |
| TFeConvNeXtLarge | Adam | **0.96** | **0.96** |
|  | RMSprop | 0.89 | 0.89 |
|  | SGD | 0.84 | 0.83 |
| TFtConvNeXtLarge | Adam | 0.94 | 0.94 |
|  | RMSprop | 0.91 | 0.91 |
|  | SGD | 0.80 | 0.76 |

**Table S3**: Performance comparison of deep transfer learning models employing ConvNextTiny under different optimizers during the 5-fold cross-validation. BAC is balanced accuracy. Best performance result is shown in bold.

| Model | Optimizer | BAC | F1 |
| --- | --- | --- | --- |
| TFeConvNeXtTiny | Adam | **0.94** | **0.94** |
|  | RMSprop | 0.91 | 0.91 |
|  | SGD | 0.80 | 0.80 |
| TFtConvNeXtTiny | Adam | 0.88 | 0.88 |
|  | RMSprop | 0.85 | 0.85 |
|  | SGD | 0.75 | 0.70 |
